# SC-Track: a robust cell tracking algorithm for generating accurate single-cell lineages from diverse cell segmentations

**DOI:** 10.1101/2023.10.03.560639

**Authors:** Chengxin Li, Shuang Shuang Xie, Jiaqi Wang, Septavera Sharvia, Kuan Yoow Chan

**Affiliations:** Department of Cardiovascular Medicine, The Second Affiliated Hospital, Zhejiang University School of Medicine, Hangzhou, 310058, P. R. China; Centre for Cellular Biology and Signalling, Zhejiang University-University of Edinburgh Institute, Zhejiang University School of Medicine, Zhejiang University, Haining, 314400, P. R. China; College of Medicine and Veterinary Medicine, The University of Edinburgh, Edinburgh, EH4 2XR, UK; Department of Computer Science, University of Hull, Hull, HU6 7RX, UK

**Keywords:** timelapse microscopy imaging, single-cell tracking, cell division, deep learning, convolutional neural networks, cell cycle

## Abstract

Computational analysis of fluorescent timelapse microscopy images at the single-cell level is a powerful approach to study cellular changes that dictate important cell fate decisions. Core to this approach is the need to generate reliable cell segmentations and classifications necessary for accurate quantitative analysis. Deep learning-based convolutional neural networks (CNNs) have emerged as a promising solution to these challenges. However, current CNNs are prone to produce noisy cell segmentations and classifications, which is a significant barrier to constructing accurate single-cell lineages. To address this, we developed a novel algorithm called Single Cell Track (SC-Track), which employs a hierarchical probabilistic cache cascade model based on biological observations of cell division and movement dynamics. Our results show that SC-Track performs better than a panel of publicly available cell trackers on a diverse set of cell segmentation types. This cell-tracking performance was achieved without any parameter adjustments, making SC-Track an excellent generalised algorithm that can maintain robust cell-tracking performance in varying cell segmentation qualities, cell morphological appearances and imaging conditions. Furthermore, SC-Track is equipped with a cell class correction function to improve the accuracy of cell classifications in multi-class cell segmentation time series. These features together make SC-Track a robust cell-tracking algorithm that works well with noisy cell instance segmentation and classification predictions from CNNs to generate accurate single-cell lineages and classifications.

## Introduction

The analysis of time-resolved fluorescent microscopy images to obtain cellular dynamics at the single-cell level has enabled the detailed study of cellular events previously inaccessible to conventional cell biological approaches [1,2]. This approach has led to the delineation of biological processes that induce a variety of cell fate decisions [3–7]. Core to this analysis is the extensive use of fluorescent markers to mark single cells and to classify cellular states. However, generating single-cell tracks from these fluorescent timelapse microscopy images is often challenging, requiring laborious optimisations of fluorescent markers and imaging conditions [2]. These optimisations are essential as good-quality cell segmentations are critical for accurate lineage tracing while avoiding biological artefacts caused by phototoxicity.

Deep learning-based convolutional neural networks (CNNs) are increasingly employed to overcome the inherent limitations of conventional fluorescence-based microscopy approaches [8]. Among the most successful applications is the use of autoencoder CNNs, enabling computationally efficient image restoration of low-light microscopy images for deconvolution, denoising and generating super-resolution image reconstructions [9]. Another area where CNNs have been successfully deployed is in the automated segmentation and classification of microscopy images [10–13]. Recent work has shown that CNNs perform well in automatically detecting, segmenting and classifying heterogeneous cellular features of microscopy images, a task previously requiring time-consuming manual human annotations [13–16].

However, applying deep learning CNNs in the automated segmentation and classification of fluorescent microscopy images presents another challenge for reliable cell tracking. State-of-the-art CNNs are inherently noisy, with instances where objects fail to be detected or are misclassified [12,13,17,18]. These inaccuracies pose a significant challenge for widely used cell tracking solutions as these algorithms are not designed to confront the various object classification errors derived from CNNs. Thus, extensive finetuning of cell tracking parameters and manual corrections of cell tracking outputs are required, posing a significant barrier for many biologists as they often lack the technical know-how and time to undertake such tasks.

To address this challenge, we developed a robust generalised cell tracking algorithm called Single Cell Track (SC-Track). SC-Track employs a hierarchical probabilistic cache-cascade model inspired by biological observations of cell division and movement dynamics of mammalian cells. We show that SC-Track can generate robust single-cell tracks from segmentation outputs from CNNs that contain missing segmentations to false detections. Furthermore, SC-Track functions well with whole cell or nuclear segmentations of diverse morphologies and imaging conditions without parameter optimisations. Finally, SC-Track can take noisy cell instance classifications and provide smoothed classification tracks to quantify cellular events accurately.

## Materials and methods

### Tracking algorithm overview

SC-Track employs a tracking-by-detection approach whereby detected cells are linked between frames. A TrackTree data structure was used to store the tracking relationships between each segmented cell temporally and spatially (Figure 1A). Each branch of the TrackTree represents a single-cell lineage of the tracked instance of a segmented cell, where branch divisions indicate cell division events and the nodes on the branches represent the segmented instances of individual cells in a specific frame. The extracted features of the segmented cell are contained in each node of the TrackTree branch.

**Fig. 1:**
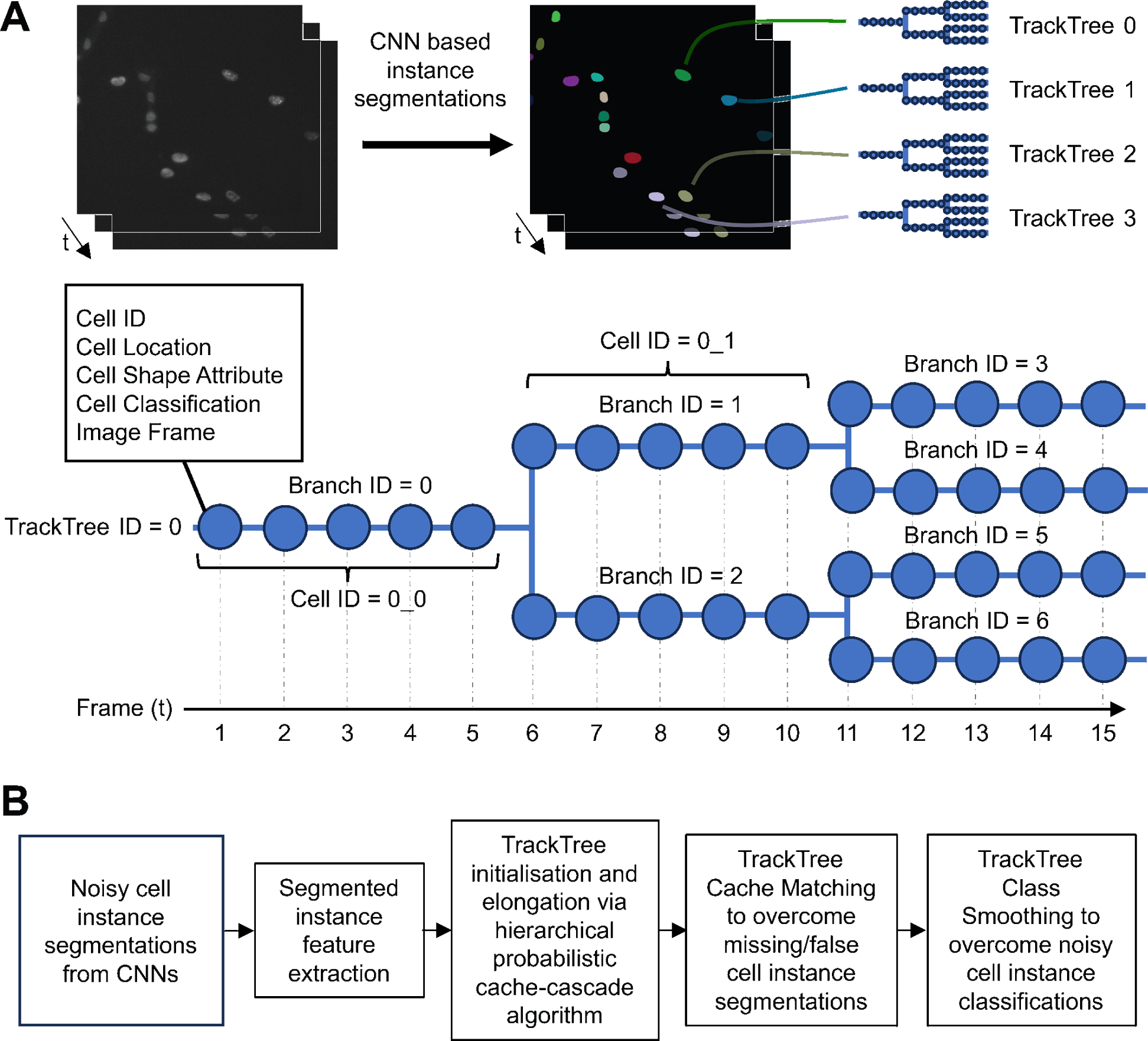
Schematic illustration providing an overview of SC-Track, the TrackTree data structure and analysis pipeline. (A) Summary of the TrackTree data structure. Each linked segmented cell is tracked in a TrackTree. A node in a TrackTree branch represents an instance of the segmented cell in a particular frame with its accompanying cell segmentation information. A branching of a TrackTree represents a cell division event. (B) Simplified overview of the analysis pipeline of SC-Track. Instance segmentation of cells from each frame is sequentially added to their respective TrackTrees. The hierarchical probabilistic cache-cascade model of SC-Track determines the assignment of each instance segmentation. If cell classification information is encoded in the TrackTrees, SC-Track will employ the TrackTree Class Smoothing (TCS) algorithm to correct the noisy cell classifications.

During the tracking process, SC-Track initialises the TrackTree list with all cells from the initial frame, representing the initial single-cell tracks for the entire time-lapse sequence. To reduce computational costs, SC-Track will attempt to connect each segmented instance with its corresponding cell from the previous frame using a hierarchical tracking approach. SC-Track will initially examine the intersection over union (IoU) of the area between segmented cells between the current frame and preceding frame (Figure 2A). Segmented cells with only one overlapping segmentation are assumed to be high-confidence links and are assigned to the corresponding TrackTree. In situations with multiple segmented cells with overlapping IoUs, SC-Track will determine the correct cells by maximising the similarity index between candidate cells between frames. When no segmented cell in the current frame overlaps with a segmented cell from the previous frame, SC-Track will expand the search area to identify possible candidates.

**Fig. 2:**
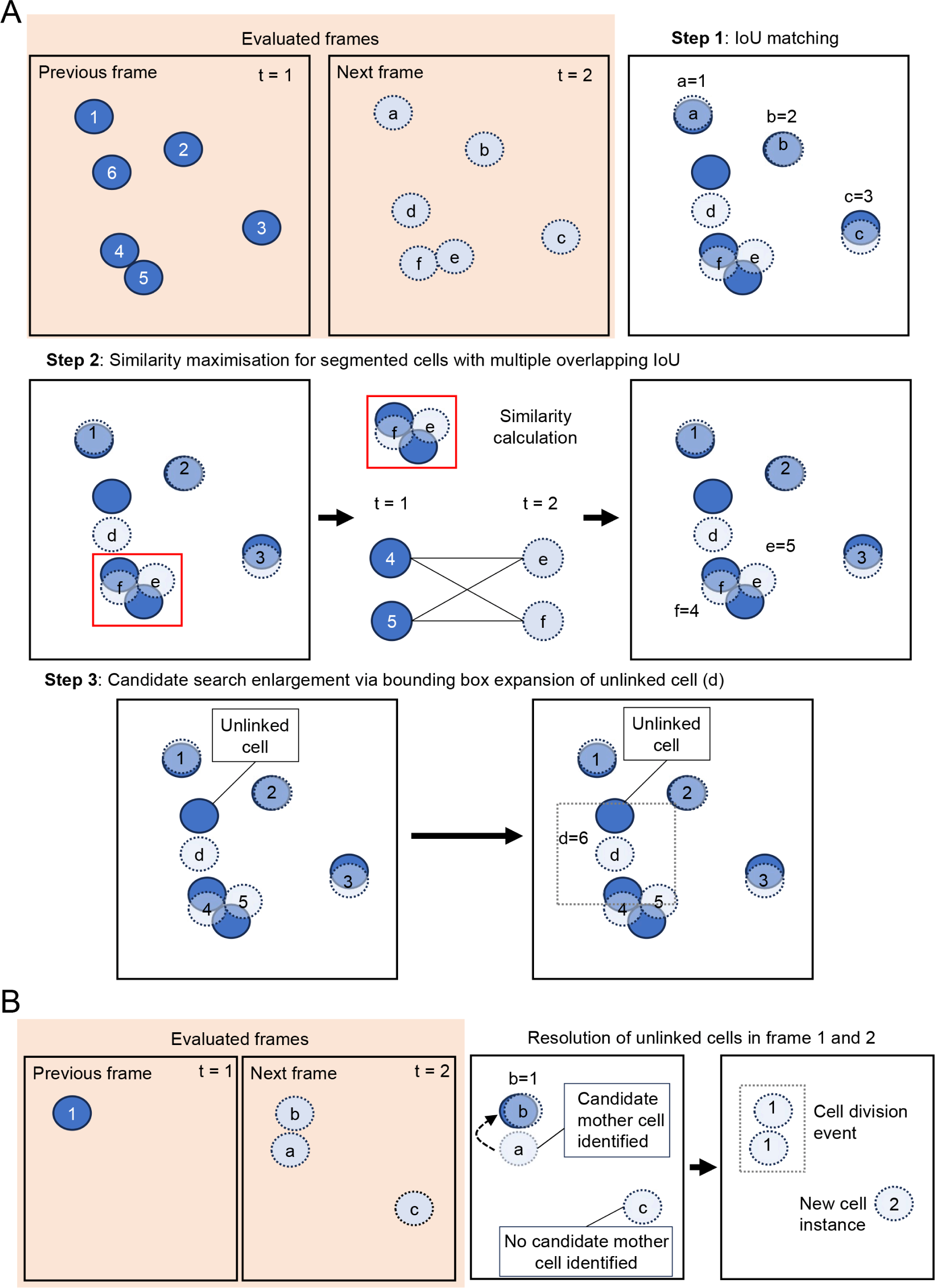
Schematic illustration summarising the hierarchical tracking approach for single-cell tracking and identifying cell division events. (A) SC-Track employs a hierarchical cell tracking approach to minimise computational costs. The overlap between the segmented cells of the preceding and subsequent frame determines the linking of segmented cells between frames. If only one cell segmentation overlaps, the segmented cell in the subsequent frame is automatically linked to the respective TrackTree in the preceding frame. When there are multiple overlapping cell segmentations, the identification of the linked cells will be determined by the similarity value of the overlapping cell segmentations. If no overlapping candidate segmented cell was identified, the bounding box of the unlinked segmented cell will be expended to identify possible candidates. (B) When a segmented cell instance that cannot be linked to available TrackTrees is identified, SC-Track will attempt to determine if a cell division event has occurred. SC-Track will perform a search in the preceding frame to determine if a compatible candidate mother cell is available. When a compatible mother cell is detected, the segmented cell instance will be branched to the corresponding TrackTree, and a cell division event is recorded. If no compatible mother cell is identified, SC-Track will assume that the segmented cell is due to a recent appearance of a cell in the microscope field of view and a new TrackTree is initialised.

By recursively searching for candidate segmented cells from the previous frame, virtually all segmented cells can be accurately assigned to the correct TrackTree. If there are more segmented cells than the number of cached TrackTrees, three possible scenarios will be considered: (1) The orphan cell is a false detection; (2) The orphan cell is an actual detection of a cell recently migrated into the field of view; (3) A cell division event has occurred leading to the generation of daughter cells.

### Detecting and assigning cell division events

In scenario three above, SC-Track will determine if a cell division occurred by searching for a potential mother cell in the mitotic state in the preceding frame (Figure 2B). When the segmented cell contains cell cycle classifications, SC-Track will allow cell division events at the TrackTree nodes where the mother cell is classified as in mitosis (M phase). However, a cell cycle-independent approach will be applied if no cell cycle information is available. In such cases, SC-Track will attempt to determine if a cell division event has occurred by matching an orphan segmented cell in the current frame to a potential mother cell in the previous frame using morphological data of the segmented cells.

To enable robust detection of cell division events without cell cycle data, SC-Track applies a series of rules based on well-established principles observed from mammalian cells undergoing cell division [19,20]. When assigning a potential mother-daughter association from a possible cell division event, the following criteria must be met: (1) At least one unlinked segmented cell was found; (2) The segmented mother cell in the previous frame must be at least 1.3 × the size of the segmented daughter cell in the following frame. (3) The candidate mother cell has not undergone a cell division event in the previous 20 frames. (4) A candidate mother cell is identified in the expanded search area of the unlinked segmented cell. If a suitable candidate mother cell was identified in the previous frame for the orphan segmented cell, the TrackTree will be branched accordingly. However, if no appropriate candidate mother cells are found, SC-Track will assume a new detection event may have occurred. The following section will discuss the algorithm employed to address the possibility of new detection events and the initialisation of new TrackTrees.

### Cache matching frames to address false and missing detection events

The stochastic nature of CNNs may lead to instances where some cells fail to be detected or are false detections [12,13,17,18], resulting in TrackTrees that contain inaccurate cell segmentations. We developed a cache-matching algorithm to address the stochastic loss of detected instances in the segmented cells (Figure 3A). In this process, SC-Track will follow a tiered response based on the likelihood that the instance segmentations are accurate. Initially, SC-Track will focus on generating high-confidence TrackTrees with cell segmentations that appear consecutively across multiple frames. When cell segmentations cannot be linked to an initialised TrackTree from the previous frame, and no suitable candidate mother cell is identified, SC-Track will search for up to five preceding frames for potential orphan TrackTrees to link (Figure 3A). If an unlinked TrackTree is found within five frames, SC-Track will assign the matched segmented cell to the corresponding TrackTree. When no matches are found, SC-Track will assume a new detection event may have occurred, and a new TrackTree will be initialised.

**Fig. 3:**
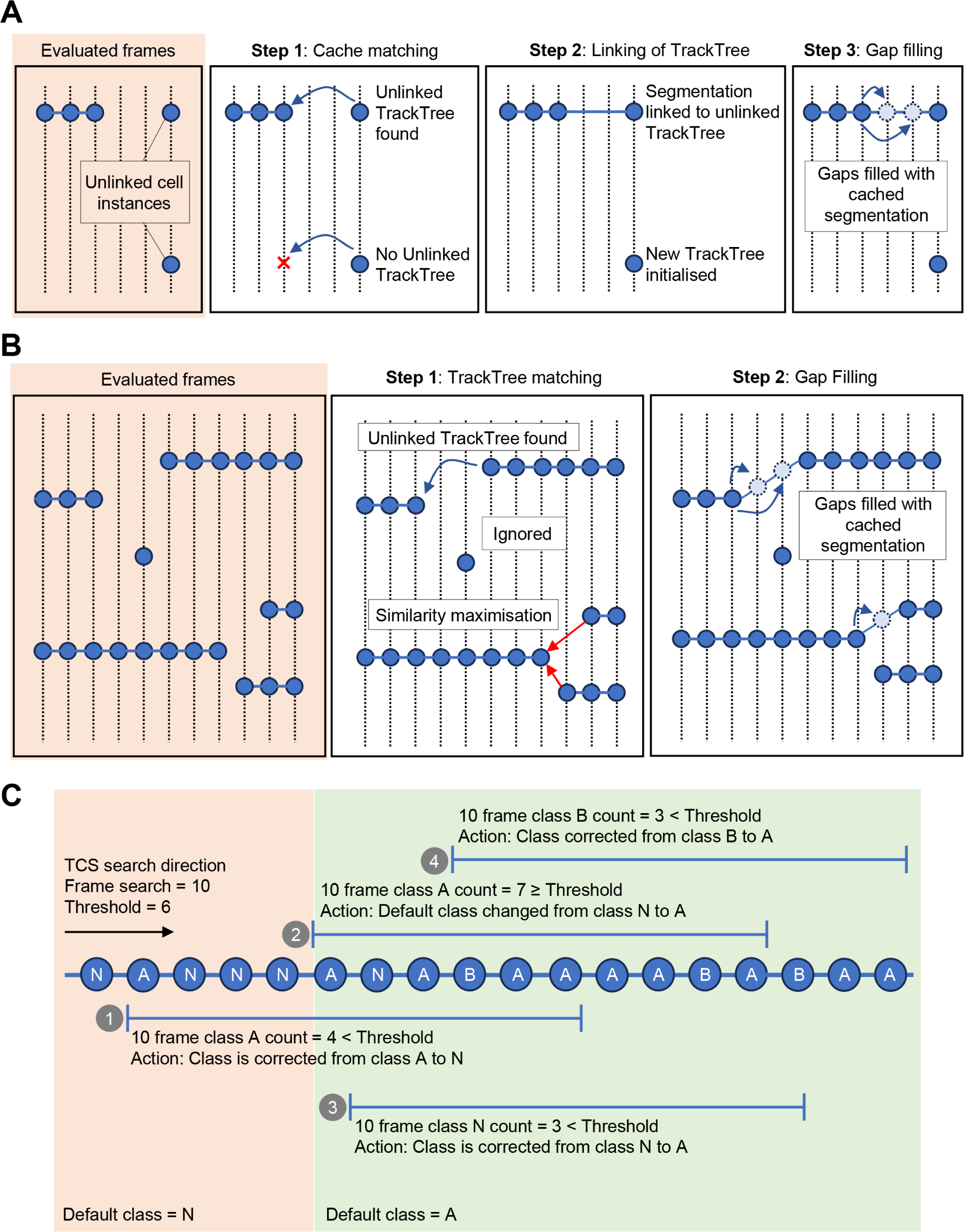
Schematic illustration describing SC-Track cached matching algorithm and TrackTree Class Smoothing (TCS). (A) Diagram describing the cache matching algorithm. When a segmented instance of a cell is detected but no compatible TrackTree is available in the previous frame, SC-Track will perform a cached search of up to five preceding frames to identify a compatible unlinked TrackTree. If a compatible TrackTree is found, the segmented instance will be linked to the TrackTree. Missing gaps in the TrackTree will be filled with the last cached cell segmentation in the preceding frame. (B) Diagram describing the TrackTree linking process. When all segmented cells are analysed, SC-Track will attempt to link TrackTrees that contain more than a single node. The linking process is based on the last segmentation of the TrackTree and the first segmentation of a separate TrackTree. When more than one compatible TrackTree is available for linkage, the cell segmentation with the highest similarity will be linked. Missing gaps in the linked TrackTrees are filled with the last cached cell segmentation in the preceding frame. (C) When a multi-class cell segmentation is performed, it is often observed that erroneous cell classifications would occur stochastically. The TCS algorithm employs a probabilistic cached search algorithm to determine if a class switch has occurred for the respective segmented cell in a time series. Note: The vertical dotted line represents a frame of a timelapse image series, and the blue circles represent an instance of a segmented cell. An established track tree is represented by a blue circle with a blue line running through the circles.

After all segmented frames are analysed, SC-Track will review all TrackTrees to filter high-confidence TrackTree initialisations (Figure 3B). SC-Track assumes that TrackTrees that contain only one node are false detections and will exclude these TrackTrees in the linking process. For TrackTrees containing at least two nodes, SC-Track will assume that some of these TrackTrees are incomplete TrackTree fragments. To identify and repair these incomplete TrackTree fragments, SC-Track will survey the terminal ends of all TrackTrees to determine instances where a terminal end of one TrackTree is spatially and temporally proximal to a terminal end of another TrackTree. When such instances are detected, SC-Track will attempt to pair the first instance of an unlinked TrackTree with the last instance of another unlinked TrackTree. The linking of TrackTrees will follow two requirements: (1) The last segmented cell of the preceding unlinked TrackTree is within the expanded search area of the first segmented cell of the unlinked TrackTree; (2) The last segmented cell of the preceding TrackTree is within three subsequent frames of the first segmented cell of the unlinked TrackTree. If more than one candidate TrackTree is available for linkage, the similarity index of the last instances of both competing candidate TrackTrees will be calculated, and the candidate TrackTree with the highest similarity will be linked. After the linking of terminal ends of track trees is completed, short TrackTree initialisations that span less than 10 frames are discarded to remove false detection instances. Finally, the intervening gaps between the linked TrackTrees will be filled with the preceding cached cell segmentation in the linked TrackTree.

### Instance classification smoothing

Instance classification of multi-class cell segmentations is often noisy [21]. We have implemented a class smoothing function to smooth out noisy classification of cells that transition from one cellular state to another. We developed the TrackTree Class Smoothing (TCS) algorithm to automatically correct the predicted results of cell type classifications (Figure 3C). TCS assumes that a cell classification change is more likely to occur in a time series when the same cell maintains the same cell classification over several frames. To evaluate the accuracy of the cell class change, TCS adopts a probabilistic cached search model. This search process is confined to the individual branch of the TrackTree and does not extend beyond the cell division branch.

The TCS probabilistic cached search model functions with the following logic: During the initialisation of the TrackTree, TCS will automatically adopt the initial classification of the detected cell instance as the default class. When TCS detects an instance where the tracked cell undergoes a cell classification change to Type A, the algorithm will undertake a forward search on the TrackTree to count the number of Type A classifications in the subsequent 9 frames. When the number of nodes classified as Type A exceeds a probability threshold of 60%, TCS will conclude that a change in cell classification has occurred and will update the default classification as Type A. Otherwise, the node where Type A was first detected will be corrected to the default cell classification. Exiting the default Type A classification occurs when a Type B class change is detected. To determine if a class change has occurred, TCS performs a forward TrackTree search in the subsequent 9 frames for the Type B classification. When the number of nodes of Type B classifications exceeds the probability threshold of 60%, the new default Type B classification is adopted. This process can be repeated to multiple cell classifications.

### Calculating similarity index when connecting segmented cells between frames

When there is more than one segmented cell overlapping with the previous frame, SC-Track will select the segmented cell with the highest similarity value with the segmented cell in the previous frame. SC-Track will employ the following formula to determine the similarity value of the possible candidate pairs between frames:

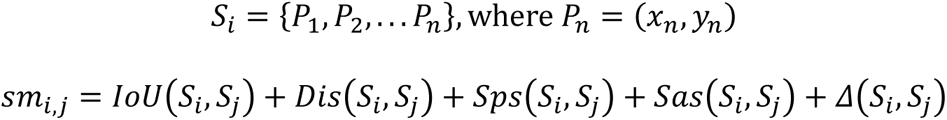

*S*_*i*_ represents the set of contour points in a 2D space defined by *x*_*n*_, *y*_*n*_ for points *P*_1→*n*_ of a cell. *sm*_i,j_ represents the similarity index between the segmented cell *i* in the previous frame and the segmented cell *j* in the subsequent frame. *Dis* is the calculated distance between the centroid of the segmented cell *i* in the previous frame and the centroid of the segmented cell *j* in the current frame. *IoU* represents the intersection over union of the contours of cells *i* and *j*. *Sps* represents the shape similarity value [22], and *Sas* represents the area similarity of the two cells. *Δ*(*S*_*i*_, *S*_*j*_) represents additional supplementary features, such as the similarity in the variance or total intensity of fluorescent signals from segmented cells. To calculate *IoU*(*S*_*i*_, *S*_*j*_), *Dis*(*S*_*i*_, *S*_*j*_), *Sps*(*S*_*i*_, *S*_*j*_), *Sas*(*S*_*i*_, *S*_*j*_), and *Δ*(*S*_*i*_, *S*_*j*_), the following formula was employed:

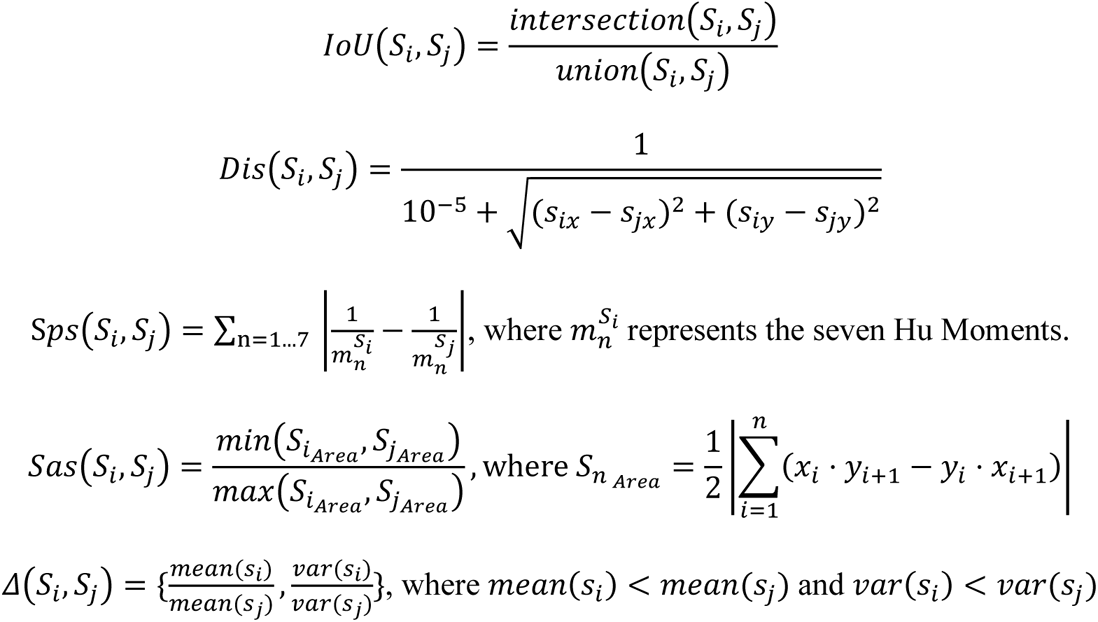

### Bounding box search area expansion

When no segmented cell in the preceding frame is detected in the segmented area of a cell in the current frame, SC-Track will expand its search area to search for potential candidates. The expansion of the search area utilises the bounding box of the segmented cell, which is expended with the following formula:

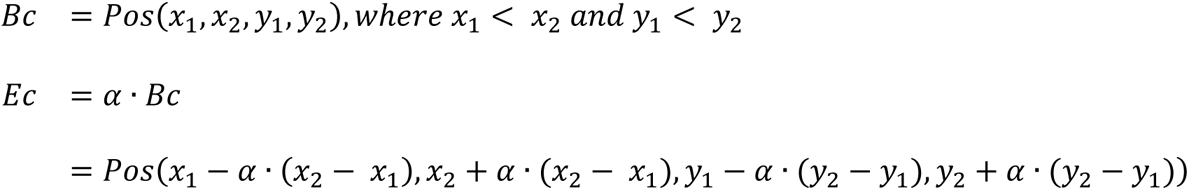

*Bc* represents the bounding box of a cell. *Pos* represents the position of the bounding box with the minimum value of the segmented cell in the x-axis and y-axis represented by *x*_1_ and *y*_1_ while the maximum value as *x*_2_ and *y*_2_respectively. *Ec* represents the expanded bounding box where potential cell candidates located in the current frame can be matched to the previous frame, *α* represents the coefficient for expanding the bounding box. By default, *α* is set to 1.

### Evaluation of cell tracking performance using MOTA and IDF1

To evaluate the performance of cell trackers in accurately tracking segmented cells, we used performance measures established in the Multiple Object Tracking (MOT) framework, which includes IDF1 [23] and MOTA [24,25]. IDF1 measures how long a tracker accurately assigns segmented cells to the correct single-cell lineage over time. It represents the ratio of correctly identified detections over the average ground-truth and computed detections [23]. IDF1 is calculated from the following formula:

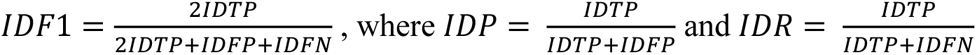

*IDP* represents the cell tracking identification precision. It is computed as the average ratio of accurately identified true positives divided by the sum of accurately identified true positives and inaccurately classified false positives. *IDR* represents the identification recall computed as the average ratio of accurately identified true positives divided by the sum of accurately identified true positives and failed detections of each single cell track.

The Multiple objects tracking accuracy (MOTA) measures the overall accuracy of the tracker performance by measuring how often a mismatch occurs between the tracking results and the ground truth [24,25]. The MOTA score is determined from the total number of errors for false positives (*FP*), missed targets (*FN*), and identity switches (*IDsw*) normalised over the total number of ground-truth (*GT*) tracks. This measure is computed using the following formula:

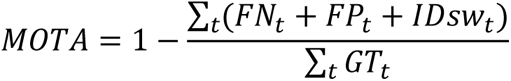

The cell tracking outputs used to calculate the IDF1 and *MOTA* values are available from Zenodo: https://zenodo.org/record/8284987. The Python scripts used to analyse the cell tracking results can be found on GitHub: https://github.com/chan-labsite/SC-Track-evaluation.

### Evaluating cell division detection events using the CDF1 score

Generating single-cell lineages over multiple cell division events requires the reliable detection of cell division events and the accurate assignment of mother-daughter relationships. We established the Cell Division *F1* score (CDF1) as IDF1 and MOTA do not measure cell division events reliably. This metric is based on the principles of the widely used *F1* score and is defined by the formula:

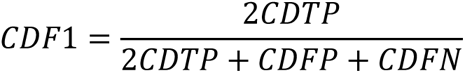

*CDTP* indicates an actual positive cell division event, where both daughter cells of a cell division event are accurately identified and assigned to the correct TrackTree. *CDFP* indicates a false positive cell division event, where daughter cells are incorrectly assigned to a TrackTree and classified as a cell division event. *CDFN* indicates a false negative cell division event, where a cell division event occurred but is not detected or the mother-daughter cells were inaccurately assigned to the wrong TrackTree. The cell tracking outputs used to benchmark the CDF1 results are available from Zenodo: https://zenodo.org/record/8284987. The Python scripts used to analyse the cell tracking results can be found on GitHub: https://github.com/chan-labsite/SC-Track-evaluation.

### Generation of datasets for cell tracking

Two cell lines with distinct morphological appearances were used to generate the imaging data to develop and test SC-Track. hTERT-RPE1 cells endogenously tagged with fluorescent mScarlet-PCNA were grown in DMEM/F-12 (Sigma, D6421) supplemented with 10% FBS (ExCell Bio, FSP500), 1× GlutaMAX (Gibco, 35050-061), 7.5% sodium bicarbonate (Sigma). MCF10A cells endogenously tagged with fluorescent mScarlet-PCNA were grown in DMEM/F-12 (Sigma, D6421) supplemented with 5% heat-inactivated horse serum (Biological Industries, 04-124-1A), 1× GlutaMAX (Gibco, 35050-061), 10 µg/ml insulin (Biological Industries, 41-975-100), 10 ng/ml cholera toxin (Sigma-Aldrich, #C-8052), 20 ng/ml EGF-β (Thermo Fisher, PHG0311), 0.5 mg/ml Hydrocortisone (MCE, HY-N0583). These cells were seeded in 8-Well chambered glass bottom slides (Cellvis, C8-1.5H-N) for two days before being imaged under a Nikon Ti2 inverted widefield fluorescence microscope equipped with a Lumencor Sola SE 365 as a light source. The cells were placed in an Okolab stage incubator (OKO) at 37°C with 5% CO_2_ and 80% humidity. The cells were observed under a 20× plan apo objective (NA 0.75), and images were captured using a Photometrics Prime BSI camera with a pixel resolution of 2048×2048. The following filter sets were used (mCherry: 560/40 nm EX, 630/75 nm EM). A single widefield image was taken in the mCherry channel (1% power, 200ms) at each stage at 5-minute intervals for up to 48 hours. A DIC image was captured at each time point (5% power, 100ms).

The timelapse microscopy images used to develop SC-Track were generated from cells cultured under the conditions described above. The images were saved as individual multi-channel TIFF files. Four time-lapse movies with varying cell densities were generated (Supplementary Table 1). These datasets were automatically segmented using a custom pre-trained StarDist model [12] and manually corrected using the VGG Image Annotator (VIA) [26] to remove false and inaccurate classifications. The annotated files contain the cell contour and “cell cycle phase” class information. The contour information was converted into a mask with values ranging from 1 to 255. The uncorrected and corrected mask images, along with the original mCherry channel image, constitute the datasets used to finetune the tracking parameters of SC-Track.

For the testing datasets, three RPE1 microscopy timelapse image series and two MCF10A microscopy timelapse image series were automatically segmented using our custom-trained StarDist model and manually corrected to ensure accuracy of the instance segmentations, cell classifications and identity of single-cell lineages (Supplementary Table 2). The imaging conditions were described above with a sampling frequency of 5 minutes. To test the reliability of SC-Track to track segmented cells with missing or false positive instances accurately, we utilised the uncorrected segmentations of the testing dataset (Supplementary Table 3). In addition, to assess how SC-Track can cope with varying levels of missing cell segmentations, we randomly deleted additional segmented cells from each frame to varying degrees to simulate higher levels of missed segmentations (Supplementary Table 4). The in-house generated segmentation masks, custom-trained StarDist model, and ground truth tracking results used in testing SC-Track can be obtained from Zenodo: https://zenodo.org/record/8284987.

### Cell Tracking Challenge dataset

We used the silver reference segmentation results from the Cell Tracking Challenge to test the SC-Track performance with various cell types and segmentation modes [27]. The silver reference datasets are derived from the uncorrected segmentation results obtained from various custom-trained CNNs applied on mammalian cell lines with different morphological appearances and imaging conditions (Supplementary Table 5). The silver reference masks and ground truth tracking results were obtained from the Cell Tracking Challenge website (http://celltrackingchallenge.net/2d-datasets/).

### Generation of single-cell lineages from segmentation masks

The segmentation results from the various evaluation datasets were used to measure the cell-tracking performance of SC-Track and three other trackers: pcnaDeep [28], Deepcell-tracking [29], and TrackMate [30,31]. For cell tracking experiments involving in-house generated testing datasets, the segmentation results in the form of a VGG image annotator (VIA2) compatible JSON file containing cell cycle class information of each segmented cell was used [26]. The data in the JSON files were read directly by SC-Track and pcnaDeep to generate the cell lineage tables. The cell segmentation data in the JSON files were converted into greyscale multi-TIFF image files before being read by TrackMate and Deepcell-tracking, as both software packages lack the function to read JSON files directly.

Default tracking settings were applied to SC-Track, pcnaDeep and Deepcell-tracking. The Lap tracker algorithm was used with default tracking settings with TrackMate. We could not perform the cell tracking experiments with the Cell Tracking Challenge dataset on pcnaDeep as it requires cell cycle class information to function [28]. The scripts used to generate the cell tracking results are available from GitHub: https://github.com/chan-labsite/SC-Track-evaluation.

### Evaluating automated cell cycle class correction

To evaluate the cell cycle class accuracy from the TCS function of SC-Track, we computed the F1 score of uncorrected raw cell classifications obtained from our custom pre-trained StarDist model and compared it with the cell cycle corrected F1 score of SC-Track (Supplementary Table 3). The F1 score is calculated with the following formula:

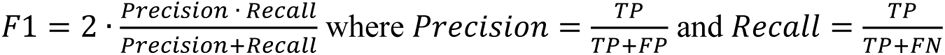

The ground truth cell cycle classification used to compute the F1 score was obtained by manually correcting the automated cell cycle classifications from our custom StarDist model. The JSON file containing the raw uncorrected cell segmentations and the cell cycle classification data used to compute the F1 results is available on Zenodo: https://zenodo.org/record/8284987. The scripts used to compute the F1 scores of individual cell cycle phases are available on GitHub: https://github.com/chan-labsite/SC-Track-evaluation.

### Runtime and multi-platform compatibility testing

We conducted runtime efficiency tests on Windows, Linux, and macOS platforms. All tests were performed using the same dataset and repeated three times. The Windows platform was equipped with an AMD R7 3700X CPU, RTX 2080 GPU, and 16GB of RAM. The Linux platform was equipped with an Intel i7 11800H CPU, RTX 3050Ti GPU, and 16GB of RAM. The macOS platform is a 2021 MacBook Pro with an M1 processor and 8GB of RAM.

## Results

### Evaluation of SC-Track cell tracking performance

The overall performance of SC-Track in cell tracking was measured using two metrics: the Multi-Object Tracking Accuracy (MOTA)[24,25] and the harmonic mean of Identification Precision and Recall (IDF1)[23]. We employed two cell tracking metrics because IDF1 is more sensitive to the total duration of incorrect track assignments, while MOTA is more sensitive to the total number of track switches [23]. We also introduced a new metric, the Cell Division F1 score (CDF1), to measure the cell tracker’s ability to reliably detect cell division events and accurately assign mother-daughter cell relationships. The CDF1 score is essential for measuring single-cell lineage reconstructions over multiple cell division events, which the IDF1 and MOTA scores cannot reliably capture. For comparison, we benchmarked SC-Track against three other freely available cell tracking algorithms that provide similar functionalities: TrackMate [30,31], Deepcell-tracking [29], and pcnaDeep [28]. The initial tests focused on generating single-cell tracks from nuclear masks obtained under ideal conditions, using manually corrected nuclear segmentation masks with accompanying cell cycle classifications with a 5-minute temporal resolution (Table 1). The results show that with ideal segmentation results, SC-Track performed best in all three measured metrics (Table 1).

**Table 1:** IDF1, MOTA and CDF1 test results based on ground truth segmentation masks.

| Dataset | SC-Track |  |  | pcnaDeep |  |  | TrackMate |  |  | Deepcell-tracking |  |  |
| --- | --- | --- | --- | --- | --- | --- | --- | --- | --- | --- | --- | --- |
|  | IDF1 | MOTA | CDF1 | IDF1 | MOTA | CDF1 | IDF1 | MOTA | CDF1 | IDF1 | MOTA | CDF1 |
| RPE1-01 | **0.994** | **1.000** | **0.965** | 0.950 | 0.957 | 0.638 | 0.902 | 0.955 | 0.308 | 0.916 | 0.983 | - |
| RPE1-02 | **0.997** | **1.000** | **0.959** | 0.957 | 0.998 | 0.533 | 0.927 | 0.975 | 0.229 | 0.935 | 0.990 | - |
| MCF10A-01 | **0.997** | **0.997** | **0.906** | 0.973 | 0.992 | 0.496 | 0.943 | 0.874 | 0.022 | 0.962 | 0.981 | - |
| MCF10A-02 | **0.982** | **0.998** | **0.971** | **0.982** | 0.978 | 0.657 | 0.945 | 0.885 | 0.000 | 0.952 | 0.967 | - |
| RPE1-03 | **0.989** | **0.994** | **0.957** | 0.975 | 0.995 | 0.308 | 0.856 | 0.923 | 0.024 | 0.931 | 0.976 | - |
| Average | **0.992** | **0.998** | **0.952** | 0.967 | 0.984 | 0.526 | 0.915 | 0.922 | 0.116 | 0.939 | 0.979 | - |

*Note: The best scores for each respective dataset and the best average score are highlighted in bold. The results for Deepcell-tracking CDF1 scores were not included as the tracker failed to detect any cell division instances in all the datasets tested.*

To further test SC-Track in generating accurate single-cell lineages, we resampled our original test dataset to mimic imaging time intervals of 10, 15 and 20 minutes. The increase in time intervals poses a more challenging cell tracking problem as each cell has more time to migrate and change morphologically between successive frames. Our results show that SC-Track gives the best overall IDF1, MOTA and CDF1 scores in the 5-minute interval, but as expected, its performance is reduced at longer intervals (Figure 4). Despite the reduced performance, SC-Track maintained the best overall performance in all three metrics across all time intervals tested. These results indicate that SC-Track works well with different temporal resolutions (Figure 4).

**Fig. 4:**
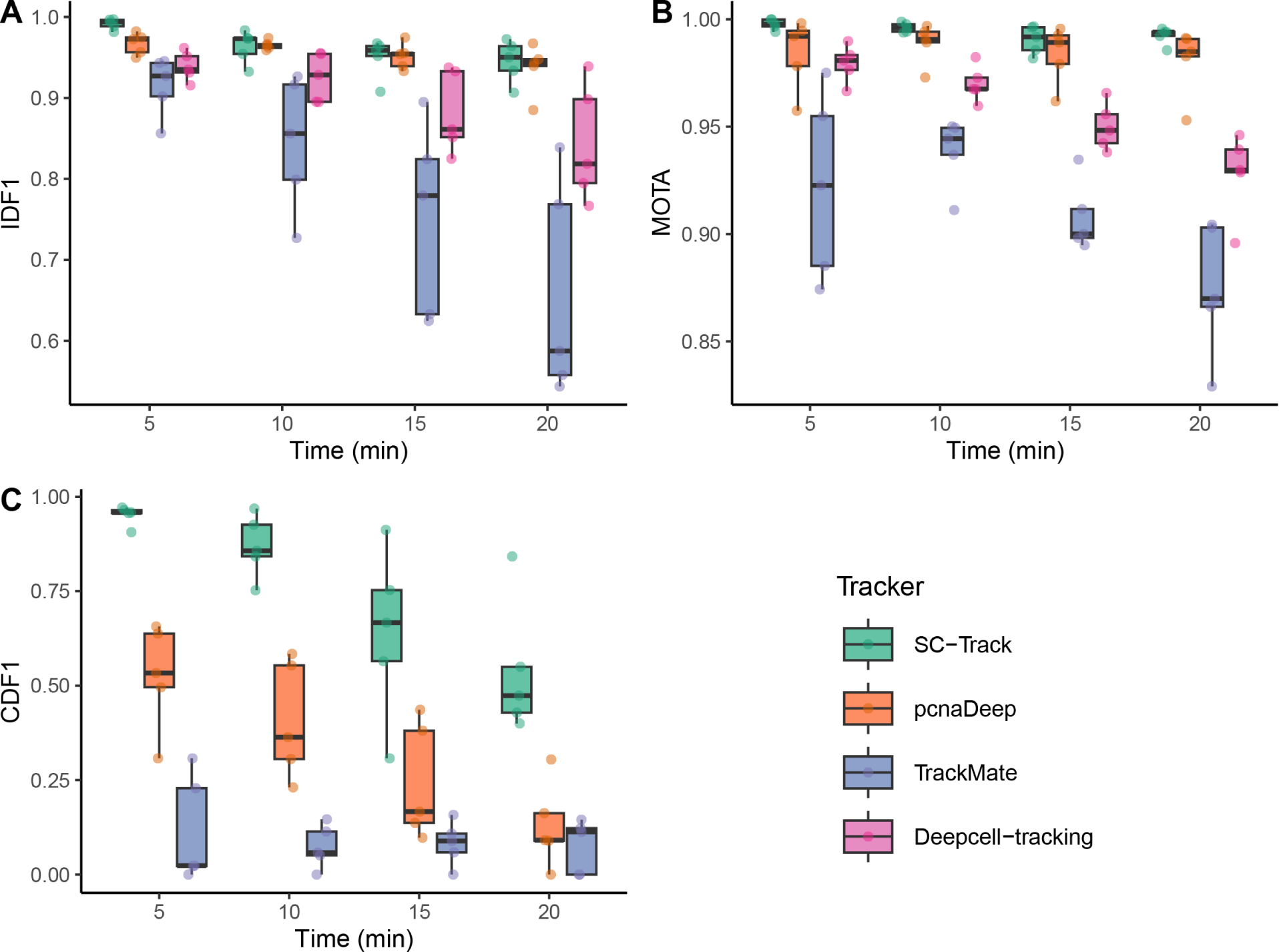
Evaluation metrics of the cell tracking accuracy based on ground truth segmentations. Box plots of (A) IDF1, (B) MOTA and (C) CDF1 scores for all four cell trackers in varying imaging time intervals. Each point displayed on the boxplots represents the respective scores of the five test datasets. The line in the boxplot represents the median. The results for Deepcell-tracking CDF1 scores were not included in (C) as the tracker failed to detect any cell division instances in all the datasets tested.

CNN instance segmentations generally display errors ranging from missing to inaccurate segmentations [12,13,17,18]. To assess the various tracker’s ability to generate reliable single-cell lineages from noisy CNN-based cell segmentations, we repeated the tests with uncorrected image segmentations from our custom-trained StarDist models (Supplementary Table 3). Our results show decreased tracking accuracy for all tested trackers, while SC-Track again gave the best overall performance, maintaining the highest average MOTA, IDF1 and CDF1 scores throughout (Table 2). To further examine SC-Track’s ability to overcome missing instances of cell segmentations, we generated a synthetic test dataset where cell instances were randomly removed at varying degrees (Supplementary Table 4). Our results show that SC-Track’s cache matching algorithm can compensate for the loss of instance detections while maintaining a high IDF1 and MOTA score throughout all tested conditions (Figure 5A and 5B). Furthermore, despite increasing missing instance detections, SC-Track remains reliable in detecting cell division events (Figure 5C).

**Fig. 5:**
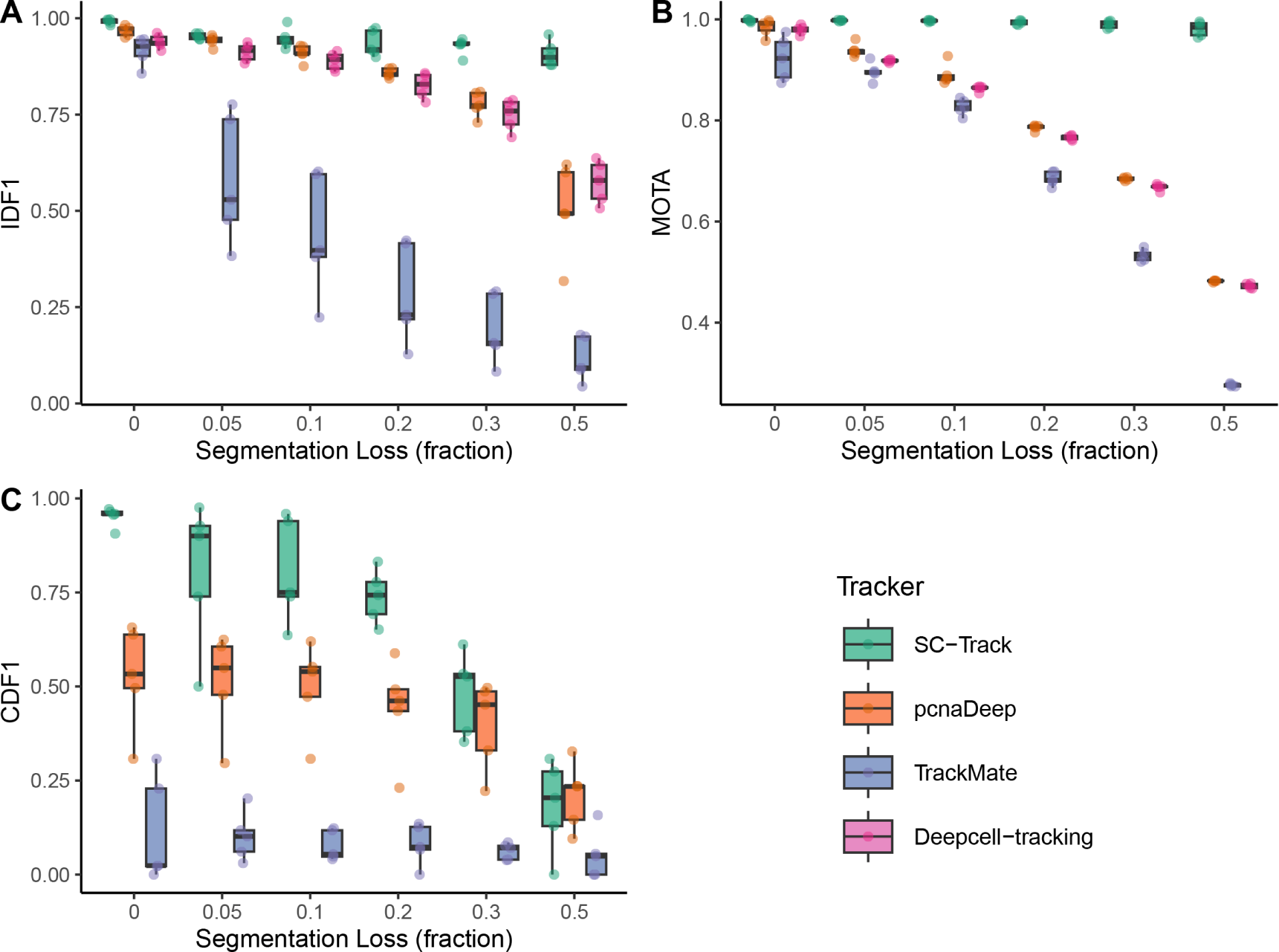
Evaluation metrics of the cell tracking accuracy based on varying cell segmentation loss. Box plots of (A) IDF1, (B) MOTA and (C) CDF1 scores for all four cell trackers challenged with the ground truth segmentation dataset with randomly deleted segmentations to mimic varying levels of cell segmentation loss. Each point displayed on the boxplots represents the respective scores of the five test datasets. The line in the boxplot represents the median. The results for Deepcell-tracking CDF1 scores were not included in (C) as the tracker failed to detect any cell division instances in all the datasets tested.

**Table 2:** IDF1, MOTA and CDF1 test results based on uncorrected segmentation masks.

| Dataset | SC-Track |  |  | pcnaDeep |  |  | TrackMate |  |  | Deepcell-tracking |  |  |
| --- | --- | --- | --- | --- | --- | --- | --- | --- | --- | --- | --- | --- |
|  | IDF1 | MOTA | CDF1 | IDF1 | MOTA | CDF1 | IDF1 | MOTA | CDF1 | IDF1 | MOTA | CDF1 |
| RPE1-01 | **0.959** | **0.996** | **0.917** | 0.953 | 0.982 | 0.391 | 0.754 | 0.876 | 0.193 | 0.929 | 0.974 | - |
| RPE1-02 | **0.939** | **0.997** | **0.768** | 0.914 | 0.980 | 0.480 | 0.746 | 0.948 | 0.088 | 0.919 | 0.971 | - |
| MCF10A-01 | **0.954** | **0.971** | **0.841** | 0.942 | 0.957 | 0.346 | 0.755 | 0.829 | 0.108 | 0.942 | 0.955 | - |
| MCF10A-02 | **0.959** | **0.978** | **0.968** | 0.944 | 0.953 | 0.286 | 0.787 | 0.874 | 0.091 | 0.948 | 0.950 | - |
| RPE1-03 | 0.964 | **0.998** | **0.952** | **0.967** | 0.995 | 0.308 | 0.771 | 0.963 | 0.081 | 0.890 | 0.958 | - |
| Average | **0.955** | **0.988** | **0.889** | 0.944 | 0.973 | 0.362 | 0.763 | 0.898 | 0.112 | 0.926 | 0.962 | - |

*Note: The best scores for each respective dataset and the best average score are highlighted in bold. The results for Deepcell-tracking CDF1 scores were not included as the tracker failed to detect any cell division instances in all the datasets tested.*

To examine the generalisability of SC-Track, we used several publicly available microscopy datasets containing diverse cell segmentations extracted from different cell types and imaging conditions (Supplementary Table 5). We decided to use the silver reference segmentation results from the Cell Tracking Challenge (CTC), as they were derived from the segmentation results of CNN models [27]. These cell segmentations are based on a collection of timelapse microscopy images taken with various imaging settings, cell morphologies and a mixture of nuclear to whole-cell segmentations [27]. Furthermore, the silver reference segmentations are accompanied by ground truth tracking results, making these datasets an impartial real-life test for SC-Track. Our results show that SC-Track consistently displayed the best cell tracking performance measured by MOTA and IDF1 scores for nearly all the CTC datasets (Table 3). Using only the silver reference segmentation results, SC-Track can reliably detect cell division events on most CTC datasets (Table 3). These results suggest that SC- Track is a general cell-tracking algorithm which performs equally well on various cell segmentation types and can be used under challenging conditions where the cell segmentation dataset exhibits high levels of detection loss.

**Table 3:** Cell Tracking Challenge imaging dataset test results for SC-Track, TrackMate and Deepcell-tracking.

| Dataset | Temporal<br>res (min) | Image | Mask | Division<br>events | SC-Track |  |  | TrackMate |  |  | Deepcell-tracking |  |  |
| --- | --- | --- | --- | --- | --- | --- | --- | --- | --- | --- | --- | --- | --- |
|  |  |  |  |  | IDF1 | MOTA | CDF1 | IDF1 | MOTA | CDF1 | IDF1 | MOTA | CDF1 |
| DIC-C2DH-HeLa | 10 | DIC | Cell | 2 | **0.992** | **0.992** | - | 0.568 | 0.639 | - | 0.206 | -0.020 | - |
| PhC-C2DH-U373 | 15 | Phase | Cell | 0 | **1.000** | **1.000** | - | 0.719 | 0.954 | - | 0.761 | 0.604 | - |
| Fluo-C2DL-MSC | 20 | Fluor | Cell | 1 | **0.979** | **0.993** | - | 0.445 | 0.685 | - | 0.588 | 0.599 | - |
| Fluo-N2DH-GOWT1 | 5 | Fluor | Nuclear | 1 | **0.991** | **0.999** | - | 0.971 | 0.948 | - | 0.902 | 0.855 | - |
| Fluo-N2DH-SIM+ | 29 | Fluor | Nuclear | 16 | **0.963** | **0.991** | **0.896** | 0.690 | 0.634 | 0.151 | 0.747 | 0.925 | - |
| PhC-C2DL-PSC | 10 | Phase | Cell | 44 | **0.989** | **1.000** | **0.977** | 0.903 | 0.945 | 0.301 | 0.879 | 0.991 | - |
| Fluo-N2DL-HeLa | 30 | Fluor | Nuclear | 56 | **0.992** | **0.998** | **0.991** | 0.881 | 0.855 | 0.061 | 0.890 | 0.968 | - |
| BF-C2DL-MuSC | 5 | BF | Cell | 5 | **1.000** | **1.000** | - | 0.143 | -0.348 | - | 0.540 | 0.096 | - |
| Average |  |  |  |  | **0.988** | **0.997** | **0.955** | 0.665 | 0.664 | 0.171 | 0.689 | 0.627 | - |

*Note: We were unable to evaluate pcnaDeep's cell tracking performance on the CTC dataset because pcnaDeep requires cell cycle data encoded in the cell segmentations to generate single cell tracks. The results for Deepcell-tracking CDF1 scores were not included as the tracker failed to detect any cell division instances in all the datasets tested. Only timelapse movies containing more than 10 cell division events are included in the analysis of CDF1 scores to ensure a reliable measure of cell division tracking performance.*

### Instance classification smoothing of single cell tracks and runtime evaluations

It is well established that CNNs occasionally misclassify multi-class instance segmentations. Instance misclassifications are caused by objects that partially fit into a particular class or suboptimal imaging conditions. The inherent noise in the cell classifications can pose a problem if accurate classifications of cellular states are essential, such as in the quantification of cell cycle phases in an image time series [28]. To overcome this problem in SC-Track, we developed a TrackTree Class Smoothing (TCS) algorithm that employs a probabilistic cached class smoothing approach to identify cell phase transition points accurately. To evaluate the utility of TCS, we measured the F1 scores of our custom-trained StarDist model to classify the cells in our test dataset using the fluorescent PCNA cell cycle reporter signal (Table 4). The results indicate that TCS can improve the F1 classification scores of all cell cycle classes.

**Table 4:** F1 cell cycle classification test results obtained from raw StarDist cell classification predictions compared with TCS corrected cell classifications.

| Dataset | Raw F1 values |  |  | TCS corrected F1 values |  |  |
| --- | --- | --- | --- | --- | --- | --- |
|  | G1/G2 | S | M | G1/G2 | S | M |
| RPE1-01 | 0.956 | 0.775 | 0.659 | **0.995** | **0.981** | **0.867** |
| RPE1-02 | 0.949 | 0.809 | 0.748 | **0.982** | **0.922** | 0.691 |
| MCF10A-01 | 0.931 | 0.690 | 0.469 | **0.982** | **0.923** | **0.580** |
| MCF10A-02 | 0.881 | 0.660 | 0.479 | **0.963** | **0.848** | **0.768** |
| RPE1-03 | 0.965 | 0.795 | 0.236 | **0.999** | **0.969** | **0.719** |
| Average | 0.936 | 0.746 | 0.518 | **0.984** | **0.929** | **0.725** |

*Note: TCS corrected F1 values that show an improvement over the Raw F1 values are highlighted in each category.*

Finally, we conducted runtime tests for SC-Track to determine how long SC-Track takes to generate single-cell tracks from cell segmentations. We measured the time taken to analyse cell segmentations from microscopy timelapse series of varying lengths (50-500 frames) and compared it with TrackMate, Deepcell-tracking, and pcnaDeep. Our results show that when working with small to moderate imaging datasets, SC-Track had the best performance (Figure 6). Increasing the number of analysed frames decreases processing speed (Figure 6), mainly because of the duration required to load the time-lapse microscopy images before generating the single-cell lineages by SC-Track.

**Fig. 6:**
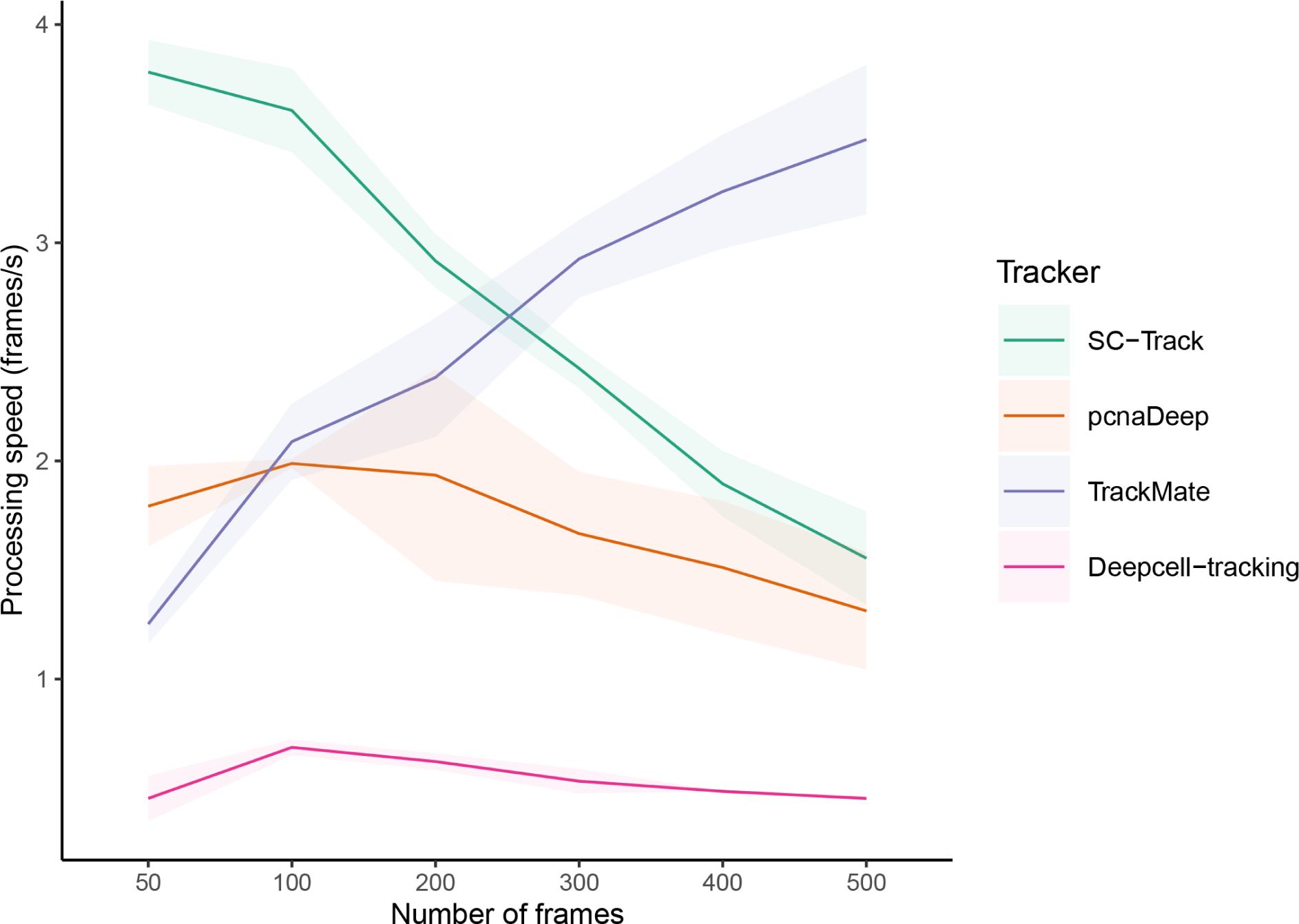
Runtime evaluation comparisons. The number of frames each tracker can process in one second is displayed on the y-axis, while the x-axis represents the varying number of processed image frames. The solid line represents the average performance, with the shaded area representing the 95% confidence interval for each cell tracker on three different computer systems running Windows, Linux, or macOS, respectively.

## Discussion

We introduced SC-Track, a novel cell-tracking algorithm employing a hierarchical probabilistic cache cascade model inspired by biologically relevant observations of cell division and movement dynamics. SC-Track can generate highly accurate single-cell tracks from nuclear and cell segmentations of diverse morphologies and imaging conditions. To better assess the ability of cell trackers to generate single-cell lineages over multiple cell division events, we introduced a new metric called the Cell Division F1 (CDF1) score. SC-Track outperformed other tested tools in detecting cell division events under all conditions tested.

SC-Track’s hierarchical probabilistic cache-cascade model tolerates false or missing cell segmentations caused by the stochastic nature of CNNs, reducing the need for extensive and time-consuming manual corrections of image segmentations. We further implemented a cache smoothing algorithm to reduce the stochastic noise in cell classifications from CNNs while increasing the accuracy of the cell classifications in time series. These functionalities are computationally efficient, allowing SC-Track to be run locally without requiring access to high-performance computing clusters.

Despite SC-Track advantages in cell tracking, there are some limitations. Firstly, SC-Track experiences a reduction in computational efficiency as the number of frames increases. This could be an issue for users who require cell tracking on high-throughput timelapse microscopy experiments. Secondly, SC-Track currently supports only 2D image analysis. Thirdly, the SC-Track class smoothing algorithm makes generalised assumptions that the degree of cell classification errors is equal across all classes and that the frequency for each cell class appearing in a timelapse series is similar. These assumptions, while reasonable, may not be valid under all biological conditions.

In summary, SC-Track solves a longstanding problem involving the use of CNNs in automated segmentation and classification of cells from time-lapse microscopy images. By encoding biological intuition in SC-Track’s cell tracking algorithm, SC-Track performs well in diverse segmentation qualities without finetuning parameters. A Python implementation of SC-Track is available, requiring only cell segmentation data and microscopy images to function, making it easy to integrate into existing analytical pipelines. These features together make SC-Track a valuable tool for biologists to help generate robust single-cell lineages and cell classifications from their timelapse microscopy images.

## Key points

- Deep learning-based convolution neural networks produce image segmentation errors that often confound well-established cell tracking algorithms.
- SC-Track is a hierarchical cache-matching algorithm inspired by biological observations of cell division and movement dynamics.
- SC-Track generates accurate single-cell lineages without parameter tuning from cell segmentations of varying qualities, morphological appearances, and imaging conditions.
- SC-Track works with features extracted from cell segmentation masks, making it easy to integrate into existing image analysis pipelines.

## Acknowledgements

The authors thank Mikael Björklund for critically reading our manuscript and for the helpful suggestions.

## Author contributions

KYC conceived the study. KYC, CL, SSX and JW conceived the experiments. CL, SSX and JW conducted the experiments. KYC, CL and SS wrote the manuscript. KYC, CL, SSX, JW and SS analysed the results. All authors reviewed the manuscript.

## Data and code availability

The raw and corrected cell segmentation results, raw cell tracking outputs, and raw analysis results are available from Zenodo: https://zenodo.org/record/8284987. The silver reference masks and ground truth tracking results are available from the Cell Tracking Challenge website (http://celltrackingchallenge.net/2d-datasets/). A Python implementation of SC-Track with its corresponding usage documentation is available at GitHub: https://github.com/chan-labsite/SC-Track. The scripts used to analyse the raw data and generate the figures presented in this manuscript are available on GitHub: https://github.com/chan-labsite/SC-Track-evaluation.

## Author Biographies

Chengxin Li is an MSc student at Zhejiang University-University of Edinburgh (ZJE) Institute at Zhejiang University, China. His research interest is bioinformatics and computer science.

Shuang Shuang Xie is a postdoctoral researcher at ZJE Institute at Zhejiang University, China. Her research interests include cancer biology, cell biology, molecular biology and biochemistry.

Jiaqi Wang is a PhD student at ZJE Institute at Zhejiang University, China. Her research interests primarily focus on cell biology, cancer, and molecular biology.

Septavera Sharvia is a lecturer at the Department of Computer Science at the University of Hull, UK. Her research interests span dependability analysis, explainable deep learning, and computational modelling.

Kuan Yoow Chan is an assistant professor at ZJE Institute at Zhejiang University, China. His research interests span computational biology, cell biology, cell cycle research and cancer biology.

---

**Supplementary Table 1:** Development dataset used in the development and finetuning of SC-Track tracking parameters.

| Dataset | Cell Type | Number of image frames | Total Cells | Average cells per frame | Temporal resolution (minutes) |
| --- | --- | --- | --- | --- | --- |
| sub-rpe-1 | RPE1 | 100 | 5374 | 53.74 | 5 |
| sub-rpe-2 | RPE1 | 100 | 6056 | 60.56 | 5 |
| sub-rpe-3 | RPE1 | 100 | 7296 | 72.96 | 5 |
| sub-rpe-4 | RPE1 | 69 | 5673 | 82.22 | 5 |

**Supplementary Table 2:** SC-Track evaluation dataset summary.

| Dataset | Original filename | Cell Type | Number of image frames | Average cells per frame | Temporal resolution (minutes) | G1/G2 | S | M | Cell division events |
| --- | --- | --- | --- | --- | --- | --- | --- | --- | --- |
| RPE1-01 | copy of 1 xy01 | RPE1 | 369 | 63.41 | 5 | 20760 | 2082 | 552 | 44 |
| RPE1-02 | copy of 1 xy19 | RPE1 | 369 | 66.16 | 5 | 22624 | 1202 | 567 | 32 |
| MCF10A-01 | MCF10A copy02 | MCF10A | 101 | 119.89 | 5 | 11147 | 541 | 345 | 29 |
| MCF10A-02 | MCF10A copy11 | MCF10A | 101 | 70.75 | 5 | 6236 | 572 | 211 | 17 |
| RPE1-03 | src06 | RPE1 | 577 | 34.50 | 5 | 18459 | 1395 | 114 | 11 |

**Supplementary Table 3:** SC-Track evaluation dataset with uncorrected raw segmentations and classifications.

| Dataset | Detected instances |  |  | Cell classifications |  |  | G1/G2 phase classification |  |  | S phase classification |  |  | Mitosis phase classification |  |  |
| --- | --- | --- | --- | --- | --- | --- | --- | --- | --- | --- | --- | --- | --- | --- | --- |
|  | TP | FP | FN | TP | FP | FN | TP | FP | FN | TP | FP | FN | TP | FP | FN |
| RPE1-01 | 23121 | 245 | 279 | 21742 | 1624 | 1658 | 19474 | 493 | 1321 | 1935 | 952 | 172 | 333 | 179 | 165 |
| RPE1-02 | 24031 | 499 | 382 | 22088 | 2442 | 2325 | 19145 | 1495 | 1190 | 2342 | 641 | 711 | 601 | 306 | 424 |
| MCF10A-01 | 11418 | 324 | 691 | 11088 | 654 | 1021 | 10008 | 418 | 756 | 562 | 35 | 124 | 518 | 201 | 141 |
| MCF10A-02 | 6812 | 120 | 334 | 5549 | 1383 | 1597 | 4818 | 414 | 1199 | 588 | 782 | 125 | 143 | 187 | 273 |
| RPE1-03 | 19651 | 158 | 255 | 18724 | 1085 | 1182 | 17392 | 366 | 916 | 1282 | 455 | 206 | 50 | 264 | 60 |

**Supplementary Table 4:**
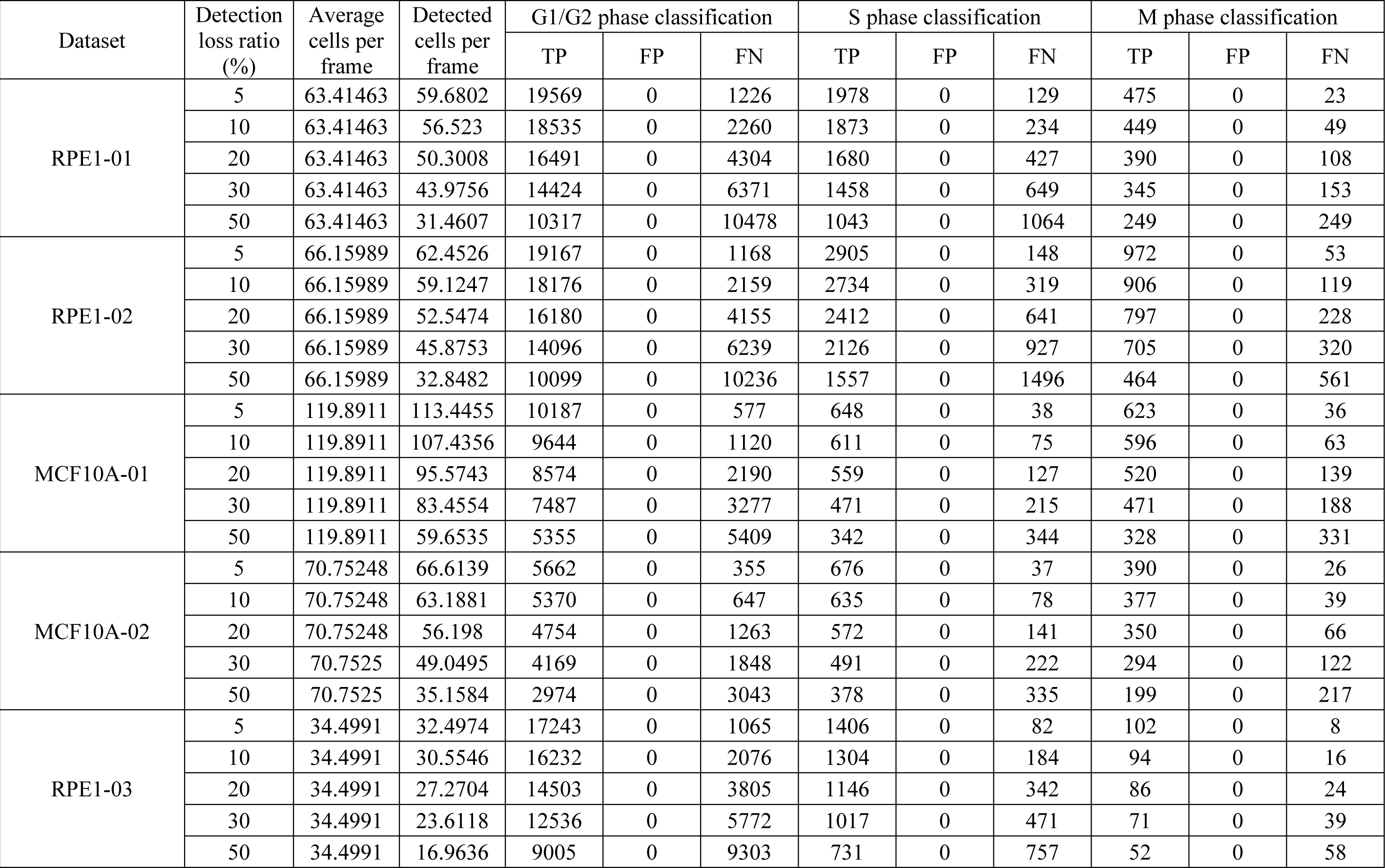
SC-Track synthetic evaluation dataset with randomly deleted segmented images.

**Supplementary Table 5:**
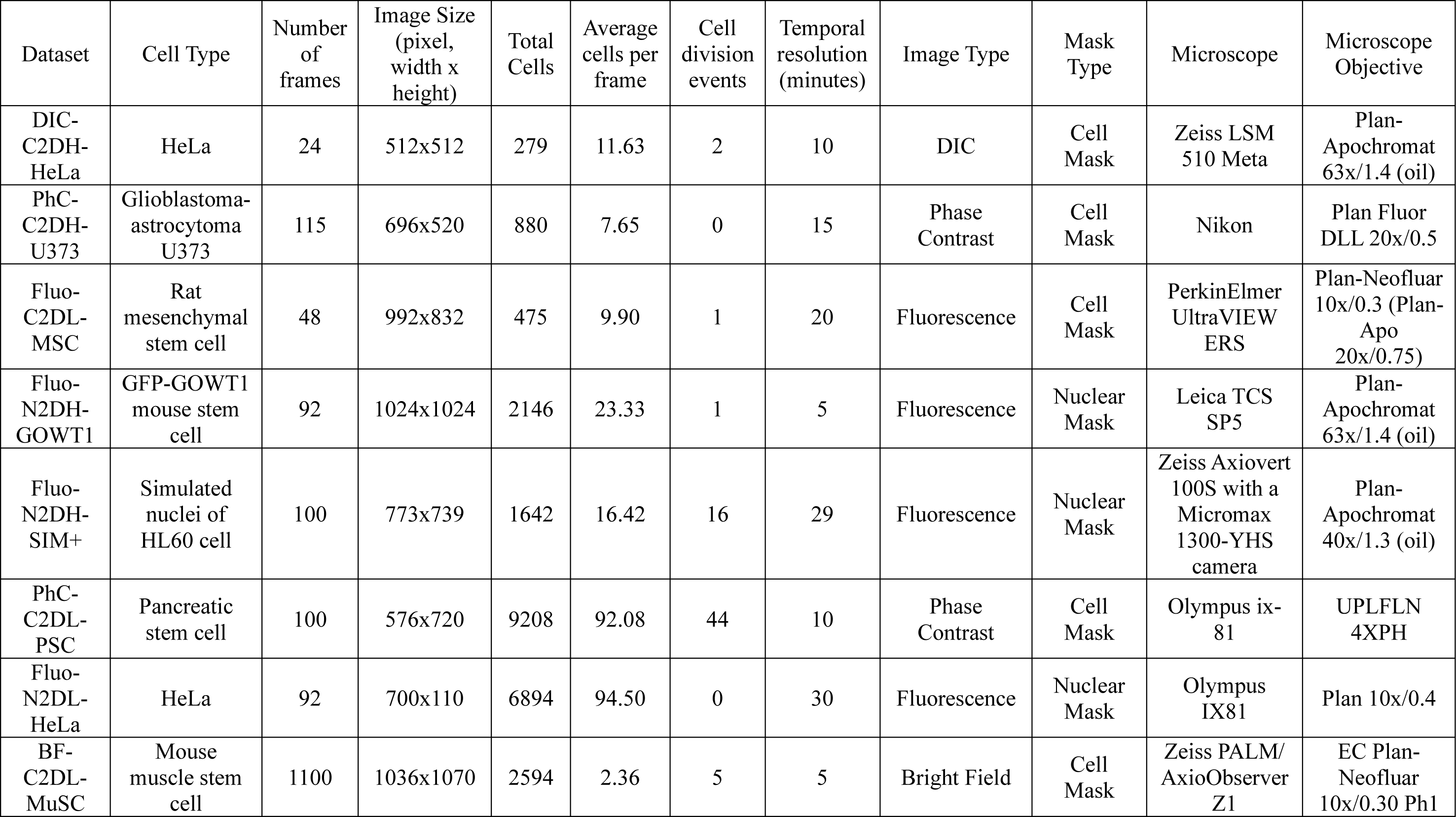
Cell Tracking Challenge Silver Reference datasets.

